# Bedside Assessment of Visual Tracking in Traumatic Brain Injury: Comparing Simple and Predictive Paradigms Using Multiple Oculomotor Markers

**DOI:** 10.1101/2025.03.27.645678

**Authors:** Shimrit Shani, Eliran Gavrieli, Oren Kadosh, Yaron Sacher, Keren Cismariu-Potash, Elena Aidinoff, Yoram Bonneh

**Affiliations:** School of Optometry and Vision Science, Bar-Ilan University; The Leslie and Susan Gonda Multidisciplinary Brain Research Center, Bar-Ilan University, Ramat Gan, Israel; Loewenstein Rehabilitation Medical Center, Department of Traumatic Brain Injury, Ra’anana, Israel; Loewenstein Rehabilitation Medical Center, Department of Intensive Care for Consciousness Rehabilitation, Ra’anana, Israel; Sackler Faculty of Medicine, Tel Aviv University, Ramat Aviv, Israel

## Abstract

Oculomotor function is a sensitive marker of neurological impairment with smooth pursuit deficiencies investigated in various disorders. However, the oculomotor deficits following traumatic brain injury (TBI) have not been fully characterized. In this study, we employed a novel bedside eye-tracking paradigm to assess oculomotor dysfunction in 30 TBI patients and 30 age-matched controls. Our paradigm utilized short, repeated linear tracking segments with head-free recording, enabling the extraction of multiple oculomotor indices, including saccadic pursuit, tracking deviation under occlusion, initial tracking speed, initial saccade latency, pupil response, and vergence instability. TBI patients exhibited widespread deficits across these indices (AUC = 0.71–0.84), which correlated significantly with functional recovery, as measured by the Functional Independence Measure (R = 0.41–0.78, p < 0.001) but not with the initial Glasgow Coma Scale scores. These findings suggest that TBI disrupts multiple components of the oculomotor system, extending to predictive tracking, pupil-linked arousal, and binocular coordination. Additionally, preliminary testing in disorders of consciousness (DOC) patients revealed fragmented tracking, suggesting a potential application for assessing perceptual awareness. Our findings support the use of eye tracking as a promising tool for quantifying brain function in TBI, with potential applications in prognosis, rehabilitation monitoring, and broader neurological assessment.

## Introduction

Traumatic Brain Injury (TBI) is one of the leading causes of death and disability worldwide; unfortunately, its incidence and associated healthcare burden are continuing to rise (Peterson and Thomas 2020). TBI often results from motor vehicle accidents, falls, or sports injuries, and it encompasses a wide spectrum of outcomes, ranging from mild cognitive disturbances to severe, life-altering impairments. Among the many deficits associated with TBI, vision and oculomotor impairments stand out as both common and disabling, with significant consequences for daily functioning (Azizi, Fielding, and Abel 2022; Debacker et al. 2018). These deficits include disruptions in smooth pursuit eye movements, saccadic accuracy, and visual tracking during occlusions—functions that rely on the intricate coordination of multiple brain regions (Barnes 2008; Keller and Heinen 1991).

Smooth pursuit eye movements allow the eyes to maintain stable fixation on moving objects by minimizing position errors. These movements involve two mechanisms: an “open-loop” phase, where the initial movement is generated without feedback, and a “closed-loop” phase, which adjusts movement based on target feedback. TBI can disrupt both phases, impairing the tracking precision and increasing reliance on compensatory catch-up saccades (De Brouwer et al. 2002; Gruber and Ahissar 2020). The sensitivity of the pursuit system to neurological disruptions makes it a valuable tool for assessing the functional impact of TBI. Predictive mechanisms also play a crucial role in smooth pursuit, particularly when tracking objects that become temporarily occluded. Previous research has shown that pursuit eye movements enhance motion prediction, consequently improving accuracy in anticipating target trajectories even when vision is disrupted (Spering et al. 2011). However, such predictive tracking can be impaired in clinical populations, including TBI patients, where deficits in motion extrapolation may contribute to tracking errors under occlusion. Such impairment was found in chronic mild TBI patients (Diwakar et al. 2015).

Several studies have examined visual tracking performance in TBI populations. For example, circular and vertical pursuit tasks have revealed slower tracking speeds, reduced pursuit gain, and more catchup saccades in TBI patients than in controls (Hunfalvay et al. 2020; Suh et al. 2006). Although these studies have demonstrated the utility of eye tracking for detecting group-level differences, they often lack correlations with clinical severity at the individual level and provide limited insight into the nuanced impairments associated with mild TBI. Furthermore, although horizontal and vertical saccadic tasks have shown promise in distinguishing TBI patients from controls (Hunfalvay et al. 2019), their applicability for mild cases or functional recovery monitoring remains largely underexplored and conceptually underdeveloped.

Visual tracking is also critical for assessing Disorders of Consciousness (DOC), where smooth pursuit often serves as a clinical marker for distinguishing Unresponsive Wakefulness syndrome (UWS) from minimally conscious states (MCS) (Trojano et al. 2012). The presence and quality of visual pursuit in DOC patients are correlated with injury severity and rehabilitation progress, making it a valuable indicator of brain function (Laureys et al. 2010). Despite its potential, assessments in this area often rely on subjective observations or stimuli that fail to account for tracking dynamics across different meridians (Candelieri et al. 2011; Giacino, Kalmar, and Whyte 2004).

In this study, we addressed these gaps by conducting a detailed analysis of visual tracking in TBI patients, focusing on smooth pursuit and tracking under occlusion. Instead of continuous circular or sinusoidal tracking used in previous studies, we employed repeated short segments of linear tracking from the center outward, allowing the extraction of multiple reliable measures. We introduced 6 novel oculomotor indices: saccadic pursuit, tracking deviation under occlusion, initial tracking speed, initial catchup latency, initial pupil response, and vergence stability, as well as assessed their correlation with clinical functional outcomes. Additionally, we applied this tracking paradigm to a subset of DOC patients to evaluate the broader applicability of these measures to assess neurological function and recovery. Our findings aim to establish eye tracking as an objective, accessible tool for TBI and DOC assessment, with implications for diagnosis, prognosis, and rehabilitation monitoring.

## Methods

### Participants

#### Main study (TBI)

The basic demographic and clinical information on the participants appears in Table 1. Initially, 33 TBI patients were recruited for the study. After excluding three participants due to poor quality eye-tracking data, the final sample consisted of 30 patients (23 severe, 3 moderate, and 7 mild) according to the Glasgow Coma Scale (GCS), who had sustained traumatic brain injury without impairment of consciousness upon entering the study. The subjects were recruited from the Levenstein Rehabilitation Hospital in Ra’anana, Israel. This study was conducted at the Loewenstein Rehabilitation Medical Center and was approved by the Helsinki Committee of the Loewenstein Hospital. The participants were aged 18-70 (M`=`39.7, SD`=`17.87), had a minimum corrected visual acuity of 6/12, and were free of existing psychiatric or neurological conditions, bilateral eye or oculomotor problems, involuntary movements of the head and body that interfere with positioning in front of a screen and eye tracking, as well as paralysis of extraocular muscles. The control group, which also included 30 people, was age-matched with the TBI group (M=40.2, SD =15.78) and shared the same inclusion criteria. All participants, including 27 men and 3 women in the TBI group, were proficient in Hebrew and underwent several tests on separate days. All TBI patients were hospitalized in the rehabilitation department for the entire study period and were evaluated by the medical staff to ensure that they could follow the instructions. Three TBI participants were excluded from the eye movement analysis due to erroneous recordings. Injury details, GCS scores, and brain imaging findings for the TBI group were obtained from medical records. GCS scores ranged from 3 to 15 (see Table 1 for detailed injury profiles). The small number of female participants reflects the male predominance of TBI cases (Siman-Tov et al. 2016). All participants provided informed consent and were told that they would not be compensated for their participation. All the experiments were conducted according to the Helsinki guidelines.

**Table 1.** Demographic data by age and gender. N = number, SD = Standard deviation, TBI = Traumatic Brain Injury, FIM1/FIM2 = Functional Independence Measure (clinical assessment at the beginning (FIM1) and around the end of (FIM2) the hospitalization. The severity level (Mild, Moderate, and Severe) was determined by the Glasgow Coma Scale (GCS).

| Group | N | Age<br>(avg ± SD) | Females/Males | FIM1<br>avg | FIM2<br>avg | GCS |
| --- | --- | --- | --- | --- | --- | --- |
| Control | 30 | 40.2 (±15.5) | 15/15 | N/A | N/A | N/A |
| TBI all | 33 | 39.7 (±17.87) | 3/30 | 56.9 | 101.9 | 6.87 (±4.75) |
| Mild | 7 | 46 (±15.49) | 2/5 | 58.8 | 102 (±18.97) | 14.6 (±0.78) |
| Moderate | 3 | 70 | 0/3 | 43.5 | 77 (±38.2) | 11 (1.4) |
| Severe | 23 | 36.1 (±16.37) | 1/22 | 60.65 | 104.34 (±20.9) | 4.17 (±1.88) |

#### DOC study

Fourteen patients were recruited from the intensive care department with a clinical diagnosis of UWS (Unresponsive Wakefulness syndrome/ Unresponsive-Wakefulness) or MCS (Minimal Consciousness State); 6 of them were excluded from the study, 2 due to a lack of cooperation, and 4 due to limited availability (one session only, which is insufficient to investigate the rehabilitation process). The patients were tested several times on separate days; some of them were available for testing for several weeks, which allowed us to assess their visual tracking improvement and their recovery from the DOC (see Table 2 in the Appendix). Five TBI patients and 1 subject with intracerebral bleeding from AVM (Arteriovenous malformation) participated in the study. The study faced challenges due to patient availability, fluctuating cooperation, and inconsistent gaze direction. Some patients deliberately avoided looking at the screen, causing us to halt the experiment.

**Table 2.**
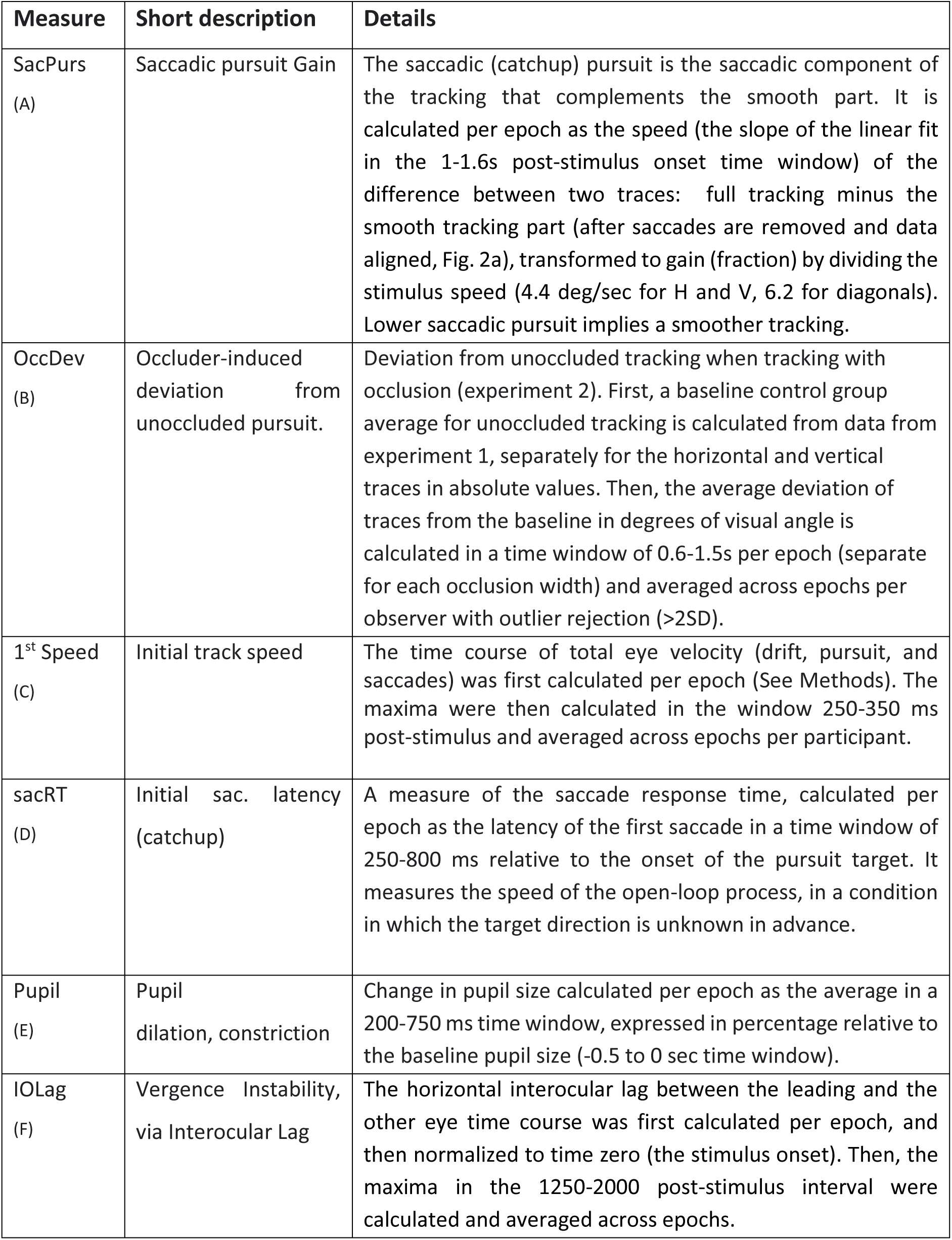
Oculomotor measures of TBI. Six oculomotor measures that reflect the quality of tracking in the two experiments are listed together with a description of their exact computation method. The measures were computed per epoch and then averaged across epochs per participant and condition with outlier rejection (see the Methods section).

### Apparatus

#### Main study (TBI)

The participants were seated in a stationary chair (without wheels) with their heads free in front of a computer display and were tested in their hospital room in its normal illumination. The stimuli were presented on a 14’’ LCD laptop monitor running in Full HD resolution (1920 x 1080) with a 60-Hz refresh rate, positioned on a standard table. A remote video-based eye-tracking system (Tobii 4c at 90 Hz sampling) was used to record eye movements and pupil size from around 60 cm, on average. The stimuli were presented binocularly using an in-house-developed platform for psychophysical and eye-tracking experiments (PSY) developed by Yoram S. Bonneh, running on a Windows PC, which was used in many of our previous studies (e.g., (Bonneh, Adini, and Polat 2015; Kadosh and Bonneh 2022; Ziv and Bonneh 2021)).

#### The DOC study

Eye tracking was performed using a remote video-based high-accuracy eye tracking system (Eyelink, SR Research, Ontario, Canada), running in a head-free mode with a 500Hz sampling rate. The testing was performed in a dedicated room in which we installed the eye tracker with a special adjustable arm on which both the monitor and the eye tracker were installed. This allowed us to adjust the display and tracker according to the position of the participant, to 50-60 cm. We built a flexible setup with which we could adjust the position of the display and eye tracker to an optimal position and used a head-free “sticker mode” for eye tracking. Although the patients did not generally move much, they were not restricted by a chinrest and some of their head movement could have affected the results. Calibration was often limited to a minimal 3-point horizontal setup or control-based methods when patient calibration failed, consequently affecting the vertical precision but not eye movement interpretation. Stimuli were displayed on a 24” LCD monitor running in Full HD resolution (1920x1080 pixels) at 100Hz running on a Windows PC, using the same presentation tool as the main study. The procedure and data analysis were similar to the TBI study; minor differences are described in the Results section.

### Stimuli and procedure

#### Stimuli

##### Experiment 1: Smooth pursuit

The stimulus sequence is depicted in Figure 1a. A fixation cross (0.47 dva) was presented on a black background for 0.5 s; then a white target disc (∼1.5 dva diameter, 35.6 cd/m² in luminance) appeared at fixation and moved from the center outwards for 1.5 s in a straight line in one of 8 directions (4 cardinal and 4 diagonal – 0, 45, 90, 135, 180, 225, 270, and 315 degrees) presented in random order. The trial sequence was repeated after a 0.5 s blank screen, with 2 trials per direction, and 16 trials per run of about 1 min. The target speed when viewed from 60cm was ∼4.4 deg/sec in the horizontal and vertical directions and ∼6.2 deg/sec on the diagonals (keeping identical horizontal and vertical speeds). Note that all space measures in degrees (degrees of visual angle, dva) were set in screen pixels and converted here to dva based on the assumed 60 cm sitting distance, which slightly varied across individuals. The stimuli for the DOC study were similar; slight changes are described in the Results section.

**Figure 1.**
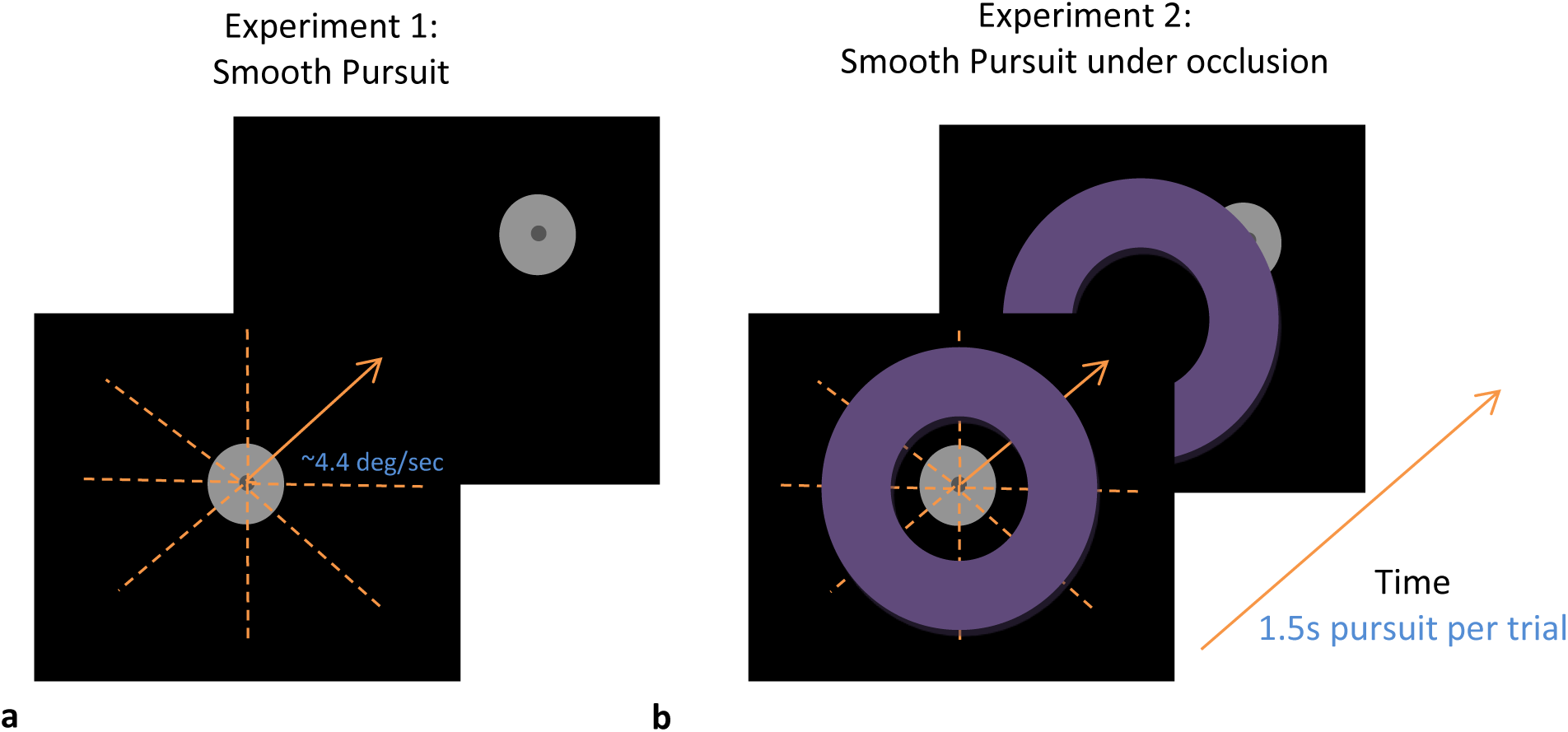
Stimuli were used for the smooth pursuit experiments. (**a**) Simple pursuit. A central fixation cross was shown for 500 ms; then a bright disk appeared and started moving smoothly in one of 8 directions in random order at ∼4.4 deg/sec speed (horizontal and vertical components are viewed from 60 cm), for 1.5s per trial, repeated 16 times per run. (**b**) Occluded pursuit. The same sequence as in (a); it was shown with the addition of an occluding ring with 3 width sizes (the medium width is shown). A third experiment was identical to the occluded pursuit experiment, except that the ring was black, i.e., invisible.

##### Experiment 2: Smooth pursuit under occlusion

The paradigm was identical to that of experiment 1, except that the target was hidden by a bright ring of varying thickness (Figure 1b). There were two types of occluder tested in separate runs: (1) A visible occluder disc with luminance of 21.6 cd/m`²`, with 3 different thicknesses (2°, ∼4°, and ∼6°) in random order, one trial per direction, 24 trials for a run of ∼1.5 minutes; (2) A virtual occluder, which was the same as the visible occluder, except that it was black as the background and therefore was invisible; its border was uncovered only when the target moved under the occluder.

##### Procedure

The nature of the study was explained to the participants, and all participants provided their informed consent to participate. The standard Tobii calibration was performed before each session. The participants were seated about 60cm from the screen. The experiments were conducted with a free head to enable hospital testing under convenient conditions including wheelchairs. Therefore, the actual sitting distance varied between observers and trials (see the Data Analysis and Results). The task was to track the moving target, without any other requirement. Each experimental run lasted for ∼1 min for experiment 1 and 1.5 min for each part of experiment 2, with a total of ∼5 min. The TBI participants performed the experiments 3 times, on average, on different days during their hospitalization, whereas the control participants did the experiments once. The clinical condition of the patients was assessed via the “Functional Independence Measure” (FIM)(Davis and Mendoza 2015; Keith et al. 1987), which is used in the LRH hospital as a clinical standard for functional evaluation. The FIM index was taken on the day of testing as the FIM closest to that day (before or after). When plotting results across days, the average of the daily FIM measures was taken. For comparison, the Glasgow Coma Scale (GCS) (Arbour et al. 2016), a standard severity measure determined during patients’ admission, was also used. In addition, dominant hand data and the direction of the head impact during the injury were collected.

### Data analysis

#### Preprocessing

Because the experiments were conducted with a free head (see Procedure), the actual viewing distance varied between observers and trials. For each trial, we calculated the viewing distance as the average distance during the interval from -0.5 s to 0 s relative to the start of tracking, excluding outliers (>2 SD). This distance ranged from 40 to 95 cm; more than 75% of the trials fell between 50 and 70 cm. Trials outside the 40–80 cm range were discarded. On average, the TBI group sat 58 cm from the screen, compared with 65 cm in the control group. The closer proximity in the TBI group was largely due to some participants using wheelchairs, which required adjustments in sitting positions for accessibility and comfort. Since target motion was defined in screen pixels, we normalized the eye-tracking data by converting them to degrees of visual angle using a fixed sitting distance of 60 cm. This ensured that data could be meaningfully compared across participants.

#### Microsaccade and blink detection

Raw traces of the two eyes’ positions and pupil sizes were analyzed to detect and remove eye blinks (along with their associated margin artifacts) as done in our previous studies (e.g., (Ziv and Bonneh 2021)), despite the difference in the device quality and sampling rate (the 90 Hz tracker *versus* the 500 Hz Eyelink). To identify blinks, we first located segments where the pupil size measurements dropped to zero. The beginning and end points of each blink were determined by examining the vertical eye position data. Specifically, we looked for deviations in the vertical trace that exceeded 4 standard deviations from the baseline, which we calculated using the mean of the initial third of our analysis window. This window extended from 100 milliseconds before to 150 milliseconds after each detected blink. Such windows outside the range of 250 to 750ms were considered missing data, which could have occurred due to head turn or movement outside the tracking range. Overall, the average missing data, together with the blinking periods, were 21%, on average, across participants and did not differ significantly between groups. For saccade detection, we used the algorithm introduced by Engbert and Kliegl (Engbert and Kliegl 2003), which is based on eye movement velocity and has been used in our previous studies (Ziv and Bonneh 2021). A velocity range of 8–400°/s and an amplitude range of 0.1–8° were allowed. Eye tracking epochs were extracted, triggered by the stimulus onset in a range of 0.5 to 3s.

#### Smooth and saccadic pursuit analysis

For calculating the “saccadic pursuit”, first, we removed the effect of saccades on the traces by erasing the saccade periods (setting to NaN in Matlab) and aligning points before and after the saccade by subtracting the displacement induced by the saccade from the trace that follows it. Then, we subtracted the traces without saccades from the full traces (that include saccades) to obtain the “saccadic pursuit” component of the traces. To compute the pursuit speed (both saccadic and smooth components), we computed linear fit estimates of pursuit speed using the Matlab statistical toolbox function “robust-fit” with outlier rejection (W < by 3SD from W mean).

#### Additional oculomotor parameters

First, we computed a set of 6 oculomotor measures described in Table 2; the exact method of calculating each measure is explained in the table. All measures were calculated per epoch and then averaged across epochs per participant and condition; outliers that exceed 2 Standard Deviations from the mean were removed, repeated twice. Group averages were calculated across participants without outlier rejection. Finally, we computed correlation R via Pearson after the above outlier rejection. We used this method of computing correlation and outlier rejection in all correlation analyses.

#### Saccade rate and eye velocity modulation functions

A saccade rate modulation function was computed to assess the event-related eye velocity modulation (Bonneh et al. 2010), following the approach used in our previous studies (Kadosh and Bonneh 2022; Yablonski et al. 2017). The effectiveness of this method can be judged by inspecting the results. The calculation was based on raw saccade onsets and proceeded as follows: for each epoch, the rate function was determined by convolving a value of 90/sec (corresponding to the sampling rate) with a Gaussian window with a sigma of 50 ms centered on the saccade onset. These rate functions were then averaged across epochs within participants, separately for each condition. To normalize the data, the participants’ mean was subtracted from each rate function, and the total average across all conditions and participants was added. Finally, the mean and standard error were recalculated across participants to derive the final modulation function. Note that the normalization process affected only the error bars and not the mean across participants. The eye velocity modulation function that quantifies movement speed including saccades drift and pursuit over time was also computed. Finally, for each epoch the total eye velocity in deg/sec was computed via a sliding window of 150 ms with a step of 22 ms, and velocity was computed as the maximal movement within the window (vector sum of the horizontal and vertical movement ranges) divided by the time window.

### Statistics

In general, we conducted statistical assessments on discrete oculomotor measures rather than on the raw traces themselves, except in two cases (Figures 2b and 4). In these instances, we analyzed the waveforms of tracking traces, pupil responses, and the interocular lag using a nonparametric permutation test (Maris and Oostenveld 2007) as applied in our previous study (Rosenzweig and Bonneh 2019). In short, for each time point, we performed a paired t-test between the two waveforms to identify the continuous significant clusters. Subsequently, we calculated the cluster-level statistics by summing all t-values within the cluster. To determine the statistical significance, we randomized the condition labels of participants’ waveforms and recalculated the group averages 1,000 times, repeating the initial step for each permutation. Finally, we computed the p-value as the proportion of permutations in which the original cluster-level t-value was exceeded by the permuted data.

**Figure 2.**
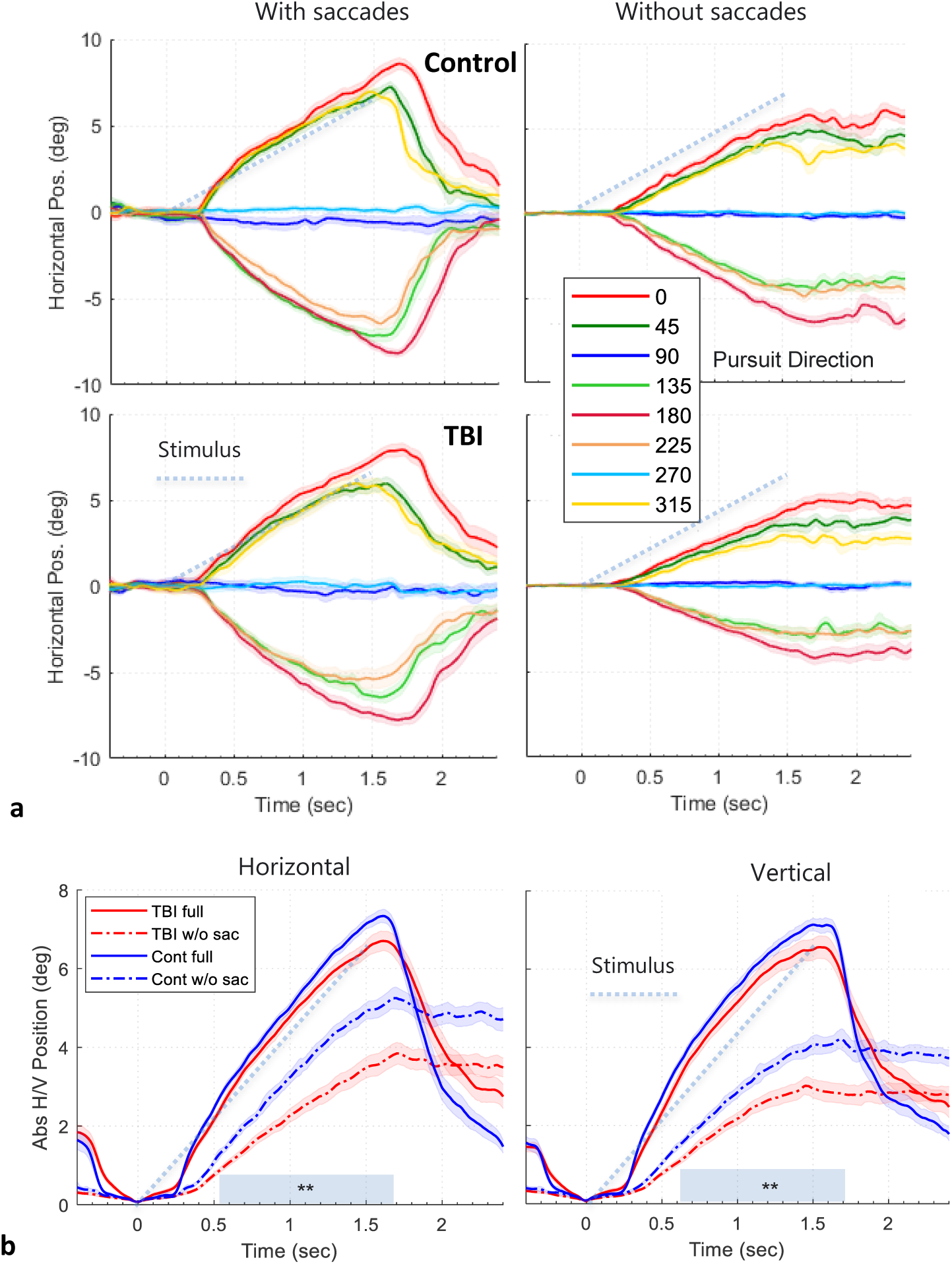
Results for smooth and catchup saccade pursuit (experiment 1). (**a**) Horizontal gaze traces as a function of time, for each target direction, normalized by the pre-trial period, averaged across trials and then across observers, for the full traces (**Left**) and for traces without saccades (after saccade removal and alignment, Right), for the TBI and Control groups (in different rows). Error bars denote 1SE across observers. Note the much shallower traces without saccades for the TBI group. (**b**) Horizontal (left) and vertical (right) gaze traces, normalized and averaged across all relevant directions (shown in (a)) in absolute values, with (solid lines) and without saccades (dashed lines) for the two groups. The smooth (without saccades) part of tracking (the dashed lines) was compared between groups via a permutation test, yielding p<0.001*** for the marked gray bar for both the horizontal and vertical traces. The light blue dotted lines show the stimulus position for the rightward and upward movement in (a), and all movements in (b).

**Figure 3.**
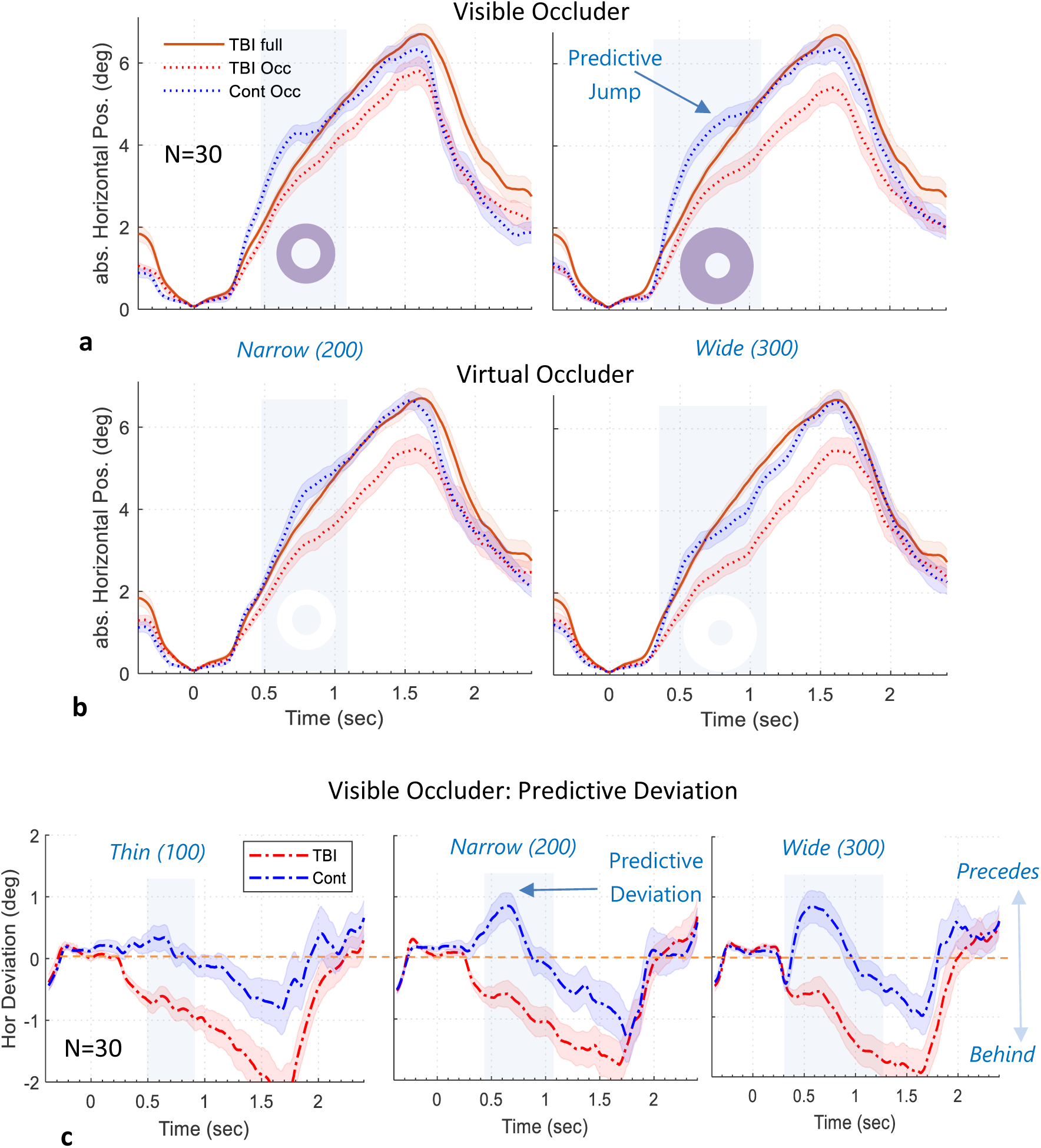
Results for smooth pursuit under occlusion (experiment 2). (**a,b**) Horizontal gaze trace averages (all directions except vertical) as a function of time, normalized by time 0, in absolute values, averaged across trials and then across observers (n=30, 1SE across subjects’ error sleeves), for the TBI (red) and Control (blue) groups in dotted curves. The red solid lines plot the TBI group average of unoccluded tracking traces from Figure 2b for reference (the Control is similar and omitted for clarity). (**a**) Visible occluder, (**b**) Virtual occluder. (**c**) The occluder-induced deviation from the unoccluded horizontal traces (all directions except vertical) for the TBI (red) and Control (blue) groups (corresponding to the difference between the occlusion traces in (a) and the unoccluded group reference, see the Methods section). Negative values correspond to lagging. Note the positive deviation of the Control group in (a) that precedes the target (especially for the wide occluder, denoted by an arrow, not found for the virtual occluder in b), whereas the TBI groups lag behind. This deviation effect is more explicitly shown in (c).

**Figure 4.**
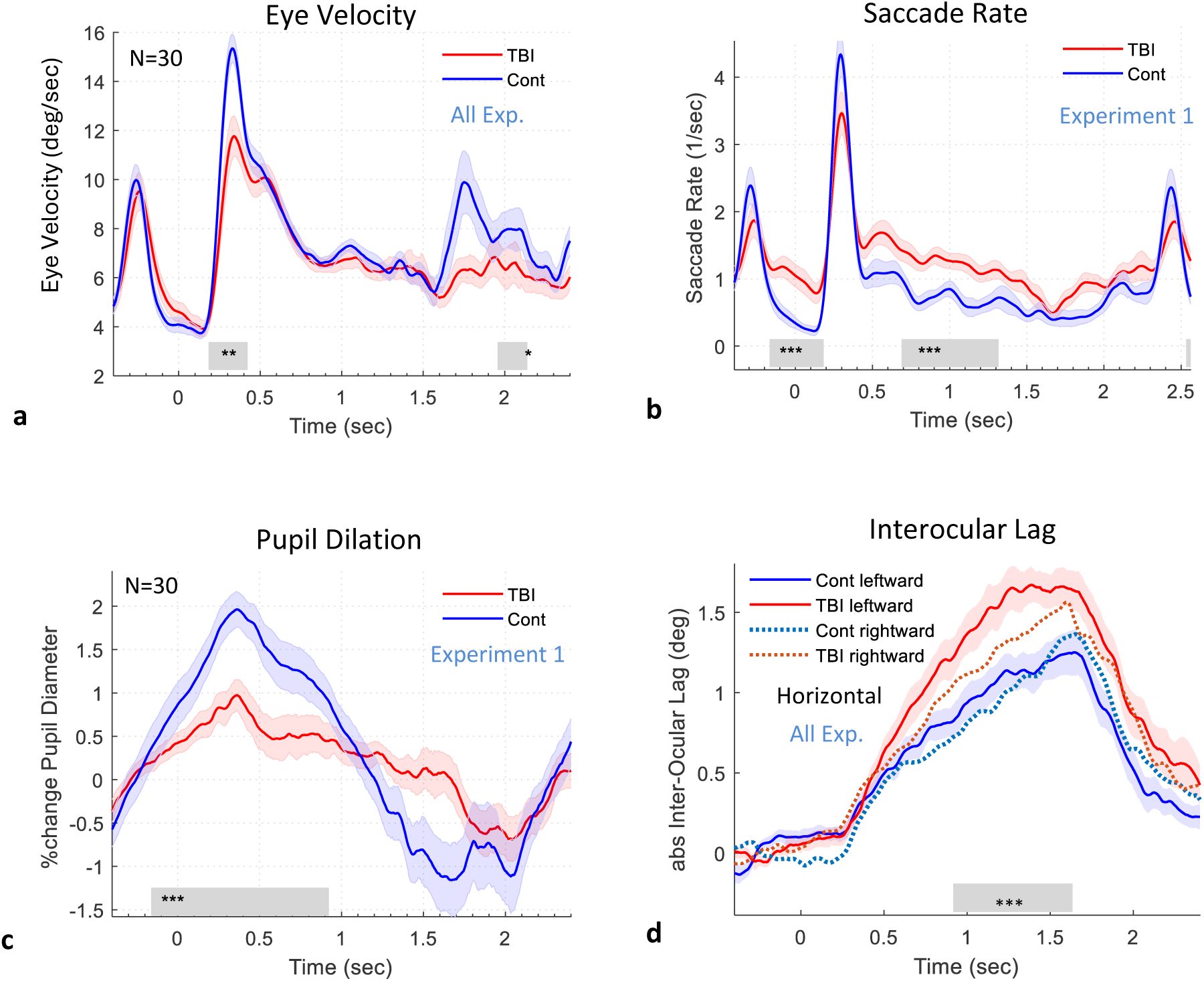
Additional tracking time-course properties, a group comparison. (**a**) Eye Velocity (including saccades, drift, and pursuit) time course averaged across all experiments. Note the faster velocity of controls around the time of the initial catchup saccade. (**b**) Saccade rate modulation in Experiment 1. Note the lower pre-trial rate in the controls, reflecting anticipatory inhibition in preparation for the stimulus onset, which is reduced in the patients. (**c**) Pupil dilation in experiment 1, reflecting the transient recruitment of arousal for starting the tracking, expressed in percentage relative to the pupil size before target onset, averaged across participants (n=30). (**d**) The horizontal interocular lag between the leading and other eyes (the gaze difference in absolute values reflecting increasing eye lead), normalized to time zero (stimulus onset) is plotted as a function of time for the horizontal tracking components, averaged across participants, separate for the leftward and rightward tracking. Only trials with a sitting distance of 50-70cm were included. Note the higher interocular lag for the patient (red), even more for leftward tracking. In all plots, the gray bars denote the period of a significant difference (the nonparametric permutation test, see the Methods section) and the error sleeves show 1SE across participants.

For computing correlations on individual participants (Figure 6, 7), first, we used the Matlab statistical toolbox function “robustfit” with outlier rejection (W < by 3SD from W mean) to extract the p-value. Then, we used the Pearson correlation to compute R with outliers excluded. For correction for multiple comparisons in Figures 5, 6, and 7, we used the False Discovery Rate (FDR) correction (Benjamini and Hochberg 1995). For assessing the group averages (Figure 5), we used ROC analysis to compute the AUC (the area under the ROC curve). Finally, we computed the effect size (the difference in SD units, aka Cohen’s D), and the p-values using the t-test.

**Figure 5.**
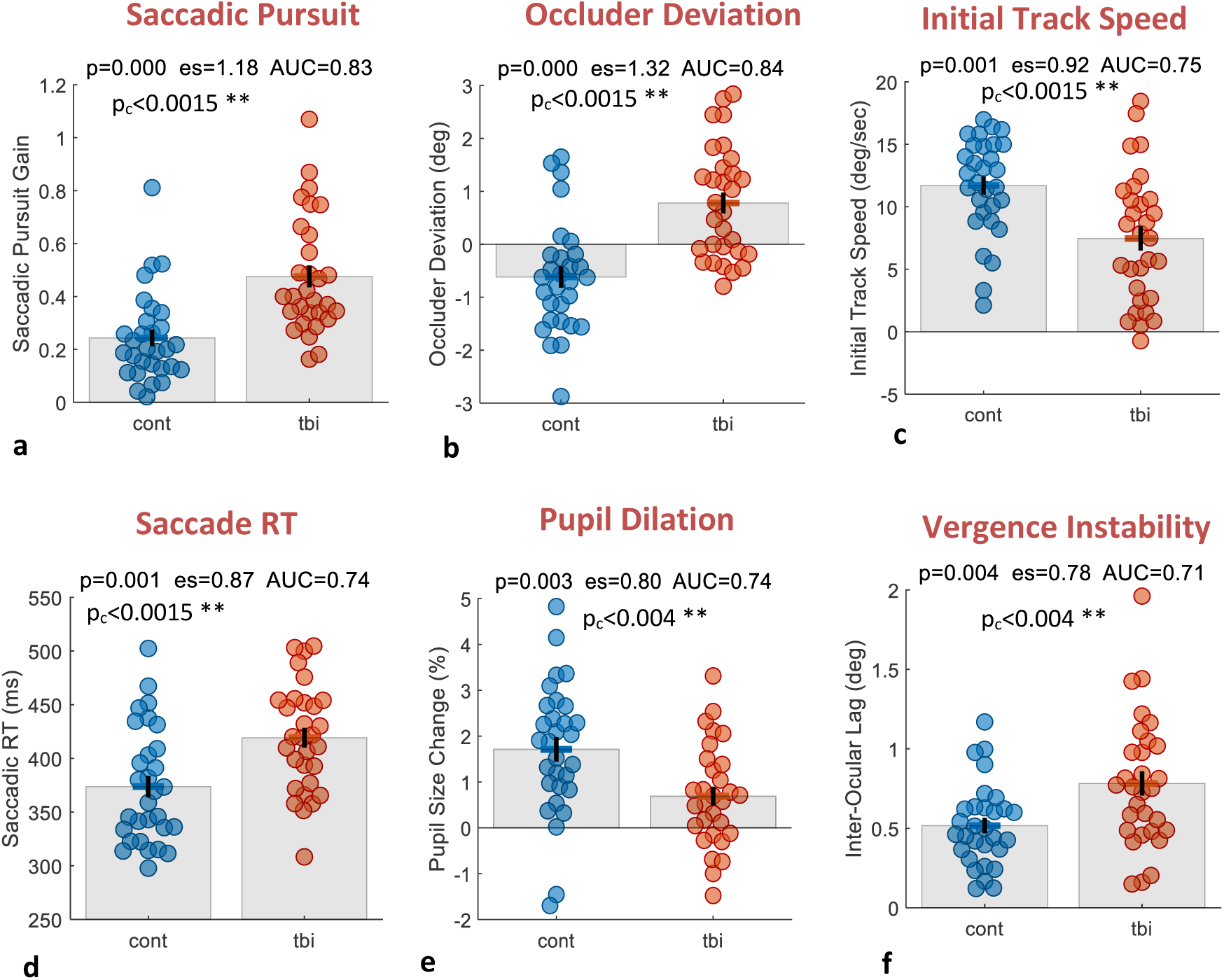
Oculomotor measures of TBI. Group differences for different oculomotor parameters (described in Table 2). For each parameter, a “bee-swarm” plot is shown with a dot per participant (N=30 per group). The error bars denote 1SE and the discrimination results are marked for each plot: p= p-value (t-test, uncorrected), p_c_=p-value FDR-corrected, es= effect size (Cohen’s D), AUC = the area under the ROC curve. As shown, all parameters showed a significant difference between groups (even when corrected for multiple (5) comparisons).

**Figure 6.**
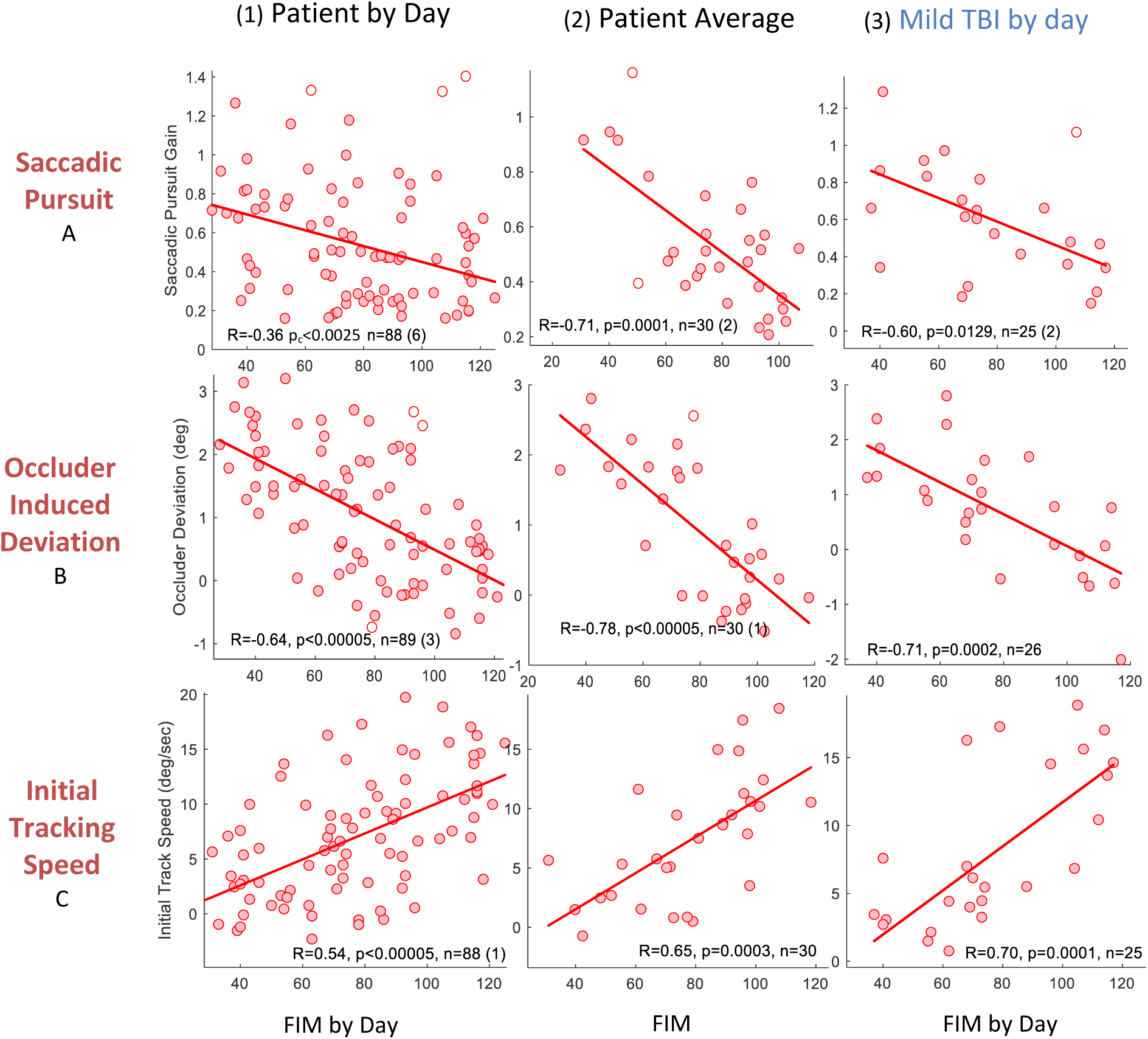
Oculomotor measures and the patients’ condition. Three measures are shown on different rows (see Table 2): (A) the Saccadic Pursuit, (B) the occluder-induced deviation, and (C) the initial tracking speed (initial “catchup”). For each parameter, three plots correlated with the Functional Independence Measure (FIM) are shown on different columns. These plots include (left to right): (1) Patient by day correlation, in which each dot corresponds to one patient on one day measured, with each patient measured 2-4 times, a total of n=∼90; (2) the patient average, with both FIM and the oculomotor measure averaged across days; (3) mild TBI patients by day, with each patient (n=7) measured 3-4 times (total of n=∼25). For each plot, the Pearson correlation R and the p-value of Robust Fit (see the Methods section) are shown at the bottom; the number of outliers rejected (if they exist) are shown in brackets, and outliers are shown in empty symbols. As shown, all correlations were significant, and most were highly significant. All p-values shown are FDR corrected for multiple comparisons.

## Results

We computed 6 oculomotor measures that we analyzed for group differences and the relation to the severity of TBI. Although there were 2 experiments, we first presented the results according to the different oculomotor measures that we extracted, some of which were accumulated across experiments (Figures 2,3,4,5). Then we analyzed the inter-relationships between measures and the cross-measures effects (Figures 6,7).

**Figure 7.**
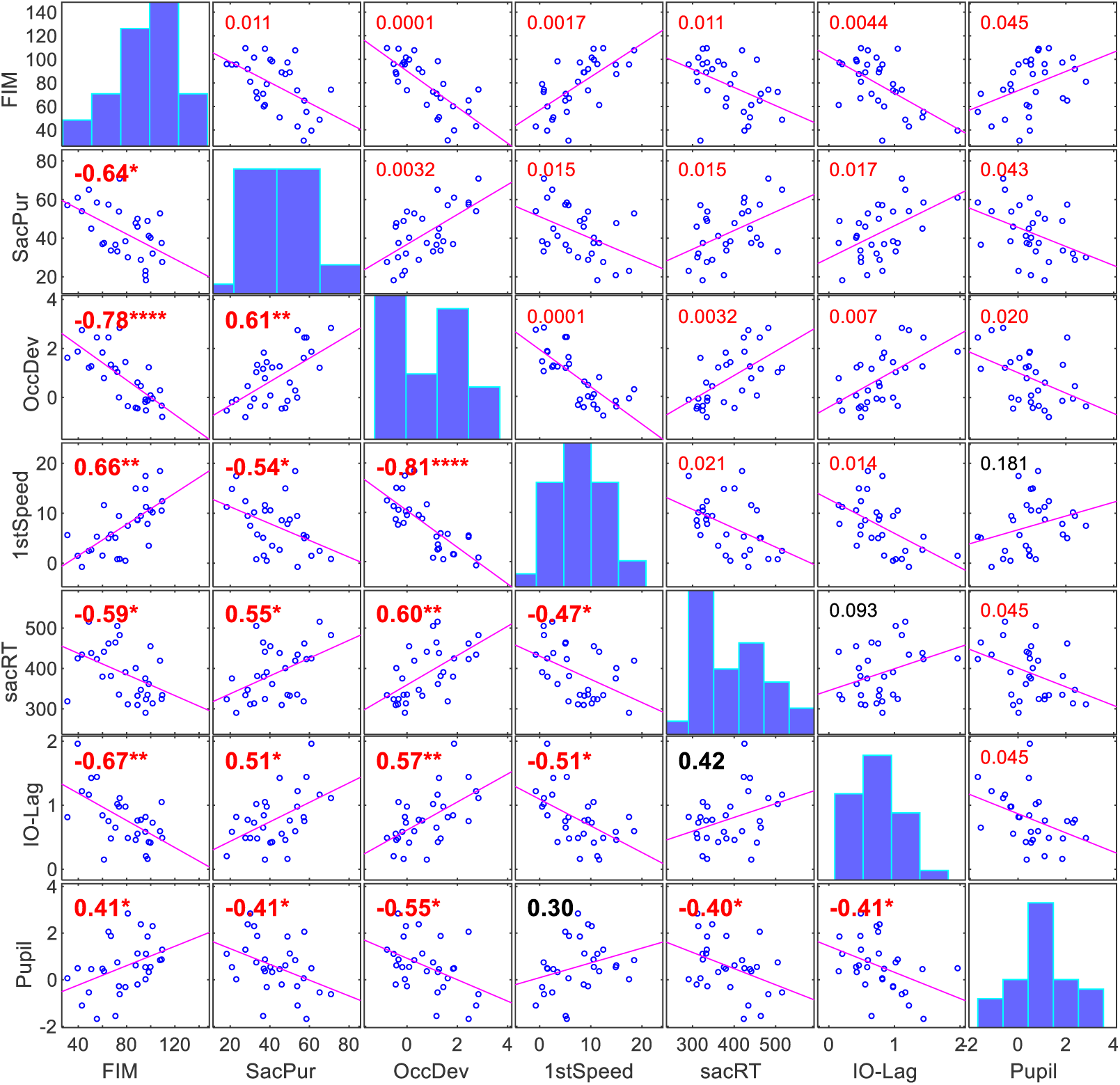
Correlation matrix of the oculomotor measures of smooth pursuit. Six oculomotor indices are correlated with each other, as well as the patient’s condition (FIM average 1,2). Each dot represents one patient. The Spearman correlation measures (R) are shown in the lower part of the correlation table, whereas the p-values (FDR-corrected for multiple comparisons, see the Methods section) are shown in the upper part. Significant correlation values are denoted in red with the standard star notation. The correlations between the patient’s condition and the oculomotor measures are shown in the leftmost column, showing that with higher (better) FIM, the TBI patients showed (1) less non-smooth “saccadic pursuit”, (2) smaller occluder-induced deviation from unoccluded tracking, (3) a shorter latency of the 1^st^ catchup-saccade, (4) less vergence instability (interocular lag), and (5) larger initial pupil dilation. All measures were significantly correlated with the clinical assessment (FIM, R=∼0.7, p<0.0005 FDR-corrected) and between each other. Age was not correlated with the patient’s condition but was modestly (significantly) correlated with the saccade RT and the occluder-induced deviation.

### Saccadic pursuit

The results of the basic smooth pursuit task (experiment 1) are shown in Figure 2. First, we plotted the group average horizontal traces for different directions (Figure 2a) for the full tracking (left column) and without catchup saccades (right column), where saccades were removed and data aligned (see the Methods section). As shown (Figure 2a left panel), both groups tracked the target similarly with a gain that approaches 1 (compared to the stimulus time course in a dotted line). To quantify this similarity, we computed the gain for each participant, averaged across trials of all directions with outlier trials rejected (outside the 0.3-3 gain range). We found an average gain of 0.88 (SD 0.14) and 0.92 (SD 0.13) for the control and TBI groups, respectively, with an insignificant difference between groups.

The results of the smooth part of tracking with traces after removing the catchup saccades are shown in Figure 2a, right column. As shown, both groups used catchup saccades, consequently making the smooth part of tracking (right column) appear “slower” in speed. However, unlike the full tracking, the smooth part of tracking was smaller in the TBI patients, as shown in Figure 2b (averages across directions in absolute values) for both the Horizontal and Vertical traces, comparing the solid lines (full tracking) and the dotted lines (smooth part, w/o saccades) between groups. We conducted a nonparametric permutation test (see the Methods section) to compare traces in the tracking time interval and found no difference between groups for the full tracking, and a highly significant difference for the smooth part of the tracking (p<0.001*** for both horizontal and vertical traces). We termed the saccadic part of the tracking “saccadic pursuit” and estimated its gain, as described in Table 2A. The results for the group comparison are shown in Figure 5a, showing a higher saccadic pursuit gain (∼0.5) for the patients compared with the controls; there was a highly significant group difference (p<0.0015 after FDR correction, an Effect Size of 1.18, and an AUC of 0.83).

### Occluder-induced deviation

An occluder, added in experiment 2, altered the horizontal and vertical traces compared with the unoccluded traces in experiment 1, causing a deviation that we assessed to determine a possible difference between groups. Three sizes of an occluding ring were tested in random order, in two types of the occluder, visible and virtual, tested in separate runs (see the Methods section). The main results of these experiments are presented in Figure 3, showing the horizontal position time course for the different directions for the two groups averaged across all non-vertical directions in absolute values, for the visible (3a) and virtual (3b) occluders. For reference, we also plotted the TBI group average of an unoccluded tracking trace from Figure 2b (the control is similar). As shown in Figure 3a, the control group “jumps ahead” when reaching the occluder, presumably a “predictive jump’, or a fast movement forward to wait for the reappearance of the target behind the occluder, upon which the tracking resumes in the typical way. The “jump ahead” effect is strongly attenuated in the Virtual condition, in which the exit point is less clear (Figure 3b). In comparison, the TBI group shows no “jump ahead” effect and resumes tracking after the occlusion, lagging, with no apparent difference between the Visible and Virtual occluders.

The difference between groups is more explicitly demonstrated in Figure 3c by plotting the occluder-induced deviation from the unoccluded horizontal traces (all directions except vertical), corresponding to the difference between the occlusion traces in Figure 3a and the unoccluded group reference (see the Methods section). Note the negative values for the TBI group, which implies lagging, and the positive deviation of the Control group that precedes the target.

To assess the occlusion effect at the group and individual level statistically, we extracted a scalar measure of “Occluder-induced Deviation” (OccDev), described in Table 2B. The results for the group comparison are shown in Figure 5b, showing a higher occluder-induced deviation from unoccluded tracking (averaged in 0.6-1.5s post-stimulus) for the patients compared to the controls; there was a highly significant group difference (p<0.0015 after FDR correction, an Effect Size of 1.32 and an AUC of 0.84).

### Initial tacking speed

We derived a discrete measure quantifying the initial tracking speed. First, we computed the time course of the eye velocity including saccades, drift, and pursuit (see the Methods section). The results, shown in Figure 4a, are pooled from all data of both the smooth tracking and occlusion experiments and from all directions. As shown, following an inhibitory period of about 150ms post-stimulus, an initial catchup movement started around ∼200 ms for both groups, peaking around ∼330ms, with a higher peak for the controls. Then, the eye velocity decreases to the tracking speed (the average of cardinal and diagonal directions, also affected by the occluder in experiment 2); then there is another higher peak around ∼1750 ms, jumping back to fixation, even more in the controls. We conducted a nonparametric permutation test (see the Methods section) that showed a significant difference around the first peak (∼200-400 ms).

To assess the initial eye velocity (shown in Figure 4a peak) at the group and individual levels statistically, we extracted a scalar measure of the “Initial Track Speed” (the 1^st^ Speed) as described in Table 2C. The results for the group comparison are shown in Figure 5c, revealing a higher initial movement (catchup) speed for the controls compared with the patients, along with a highly significant group difference (p<0.0015 after FDR correction, an Effect Size of 0.92 and an AUC of 0.75).

### Saccade RT and inhibition

First, we analyzed the saccade rate modulation (see the Methods section); the results are shown in Figure 4b. As shown, the control group inhibited the saccades before the onset of the target, reflecting temporal anticipation, which was much weaker in the patients. To assess the significance of this effect, we conducted a nonparametric permutation test (see the Methods section), which revealed a significant difference (***) in the pre-stimulus period and immediately after (∼-200-200 ms, Figure 4b). There was also a highly significant difference during the tracking period, consistent with the “saccadic pursuit” measured, described above.

We conducted an additional analysis of the latency of the initial catchup saccade, which we termed “Saccade RT”, described in Table 2D. Since in all experiments the target appeared at fixation and started moving in one of 8 directions unknown to the observer, the measure of Saccade RT represents the processing speed of identifying the direction of the target and starting the tracking (the open-loop stage). The group comparison for data from all experiments is shown in Figure 5d, showing an earlier initial saccade for the controls compared to the patients (374 *vs.* 420 ms), with a highly significant group difference (p<0.0015 after FDR correction, an Effect Size of 0.87, and an AUC of 0.74). Note that although in the saccade rate modulation function (Figure 4b) there is no apparent difference in the initial peak (around 300 ms), the saccade RT is based on the latencies of the initial saccades in an interval of 250-850 ms and many saccades were delayed in the patients.

### Pupil Dilation

The initial catchup saccade was found to be associated with dilation of the pupil, which reflects recruitment of arousal to drive the overt attention shift (Stephanie Jainta, Marine Vernet 2011). First, we analyzed the pupil size modulation time course for the two groups; the results are shown in Figure 4c. As shown, the pupil dilated relative to the pre-stimulus baseline, peaking around 350ms post-stimulus, with ∼2% dilation for the controls and only ∼1% for the patients, an effect found highly significant (nonparametric permutation test, see the Methods section). Next, we extracted a scalar measure of the Pupil Dilation, as described in Table 2E. The results for the group comparison are shown in Figure 5e, showing a higher pupil dilation for the controls compared with the patients, with a highly significant group difference (p<0.004 after FDR correction, an Effect Size of 0.8, and an AUC of 0.74).

### Interocular Lag

We observed that when the eyes start to track a target moving from fixation horizontally, they start deviating, compared with the pre-tracking time, with increasing interocular lag over time up to 1-2 deg (Figure 4d). We quantified this effect by subtracting the horizontal gaze position of the two eyes to compute the interocular lag, normalizing it to the interocular difference at time zero, and taking the absolute value to avoid the effect that the eye leads. In this way, we focused only on the deviation induced by the tracking. Note that for this analysis, we considered the individual sitting distance when converting the traces to degrees of visual angle (dva), although the results persist without this correction. The results are shown in Figures 4d and 5f, with data from experiment 1 (simple tracking). As shown, there was a larger interocular lag in the patients compared with controls in the horizontal component of tracking, and this lag increased during the tracking. We quantified it via the discrete Interocular Lag measure (Table 2F); group averages are compared in Figure 5f, showing a significantly larger interocular lag in the TBI group (p<0.004 after FDR correction, an Effect size of 0.78 and an AUC of 0.71). There was also an interocular lag on the vertical movement, but this effect was much smaller and did not reach group significance.

### Comparing the oculomotor measures and the clinical condition

To assess the relationship between the oculomotor measures and the clinical assessment of the patients, we first considered the two clinically available measures, (1) the GCS, which was determined at the time of admission and does not reflect the current condition of the patient, and (2) the FIM (Functional Independence Measure, see the Methods section), which could change over time and was assigned per patient and on the day of testing.

First, we examined three measures in more detail: (1) the Saccadic Pursuit, (2) the Occluder-Induced Deviation, and (3) the Initial Tracking Speed, which we correlated with the FIM in three ways: (a) a measure per subject and day, resulting in ∼90 measures, (b) a measure per subject, and (c) a measure per subject (n=7) and day (3-4) for the Mild TBI (mTBI) patients selected by GCS>=13. The results are shown in Figure 6. As shown, all correlations were significant, and most are highly significant, even after FDR correction (see the Methods section). The results indicate the applicability of these measures to assess the patient’s condition, including the changing condition of the patient, for patients of varied severity, including those considered “mild” or mTBI due to the initial classification.

To obtain a more complete picture of the relationships between the oculomotor measures and the patients’ condition, we computed a correlation matrix for the patient data that reveals the inter-relationship and consistency between the measures as well as the clinical assessment and the age of the patient. The results are shown in Figure 7. As shown, the different oculomotor measures were remarkably correlated with the average FIM (assigned per subject and on the day of measure), with absolute R in the range ∼0.6-0.8, except for the pupil (R=0.41) and p-values (after FDR correction) between 0.0001 and 0.045. The highest correlation was found for the Occluder-Induced Deviation (R=0.78). In all measures, the direction of correlation was consistent with the group average comparison to controls (Figure 5), i.e., higher (better) FIM was associated with oculomotor measures that were similar to the healthy controls. All measures were mostly significantly correlated with each other, but with a lower correlation, indicating that these measures were not redundant.

The sitting distance, computed per trial at the start of tracking (see the Methods section), varied both across participants and within participants (due to free head movement). We excluded two control participants because they sat too far from the screen (an average distance >85 cm). To assess whether the sitting distance influenced our findings, we computed correlations between each participant’s average sitting distance and the parameters analyzed in Figure 5. There were no significant correlations, nor any indication of a trend, with p-values ranging from 0.2 to 0.9 in the TBI group and 0.5 to 0.7 in the Control group. Similarly, we found no significant correlation, nor any trend between the sitting distance and the individual average Functional Independence Measure (FIM) scores. These findings confirm that the observed differences between the TBI and control groups—and among patients themselves— cannot be attributed to variations in the sitting distance, thereby supporting the validity of our results. We also examined the correlation of the same measures with the GCS and found no significant correlation or trend for any of them (p>0.5 for 4 of the measures, p>0.15 for the pupil and msRT).

### Pursuit eye movements in DOC

A separate part of the project was devoted to testing patients with a Disorder of Consciousness (DOC) on the same tracking paradigm but with slight differences. This study was performed in the department for the rehabilitation of Consciousness at Lowenstein Hospital with 11 participants (see Table S1 and Table S2); however, 6 of them were excluded due to unavailability or lack of cooperation. See more details in the Methods section.

The individual horizontal tracking results for the patients and a group average of control subjects (n=5) are shown in Figure 8, with data averaged across all testing days. The results dissociate between saccadic pursuit and smooth pursuit (saccades removed and data aligned) for left and rightward movement separately. The results are described in detail in the figure caption. The results reveal varied tracking performance across patients. For example, patient MCS 5 exhibited almost a normal tracking amplitude with saccades, whereas patient MCS1 exhibited almost no tracking at all; 3 other patients exhibited different degrees of intermediate tracking amplitude (MCS2-MCS4). The results also reveal a striking difference between the full (saccadic) tracking and the smooth pursuit part of it. Whereas saccadic tracking was significant and sometimes almost normal (MCS5), the smooth pursuit component was very small if not negligible. For example, patient MCS5 showed a smooth pursuit speed equivalent to 15% of the saccadic pursuit speed, with similar percentages in the other patients. This shows that the patients could not perform closed-loop tracking and were compensated with open-loop catchup saccades. Another interesting result is the asymmetry between leftward and rightward tracking found in 2 of the patients (MCS2 and MCS4). Patient MCS2 exhibited a leftward tracking amplitude of 30% of the rightward tracking amplitude, which was by itself 50% of normal tracking.

**Figure 8.**
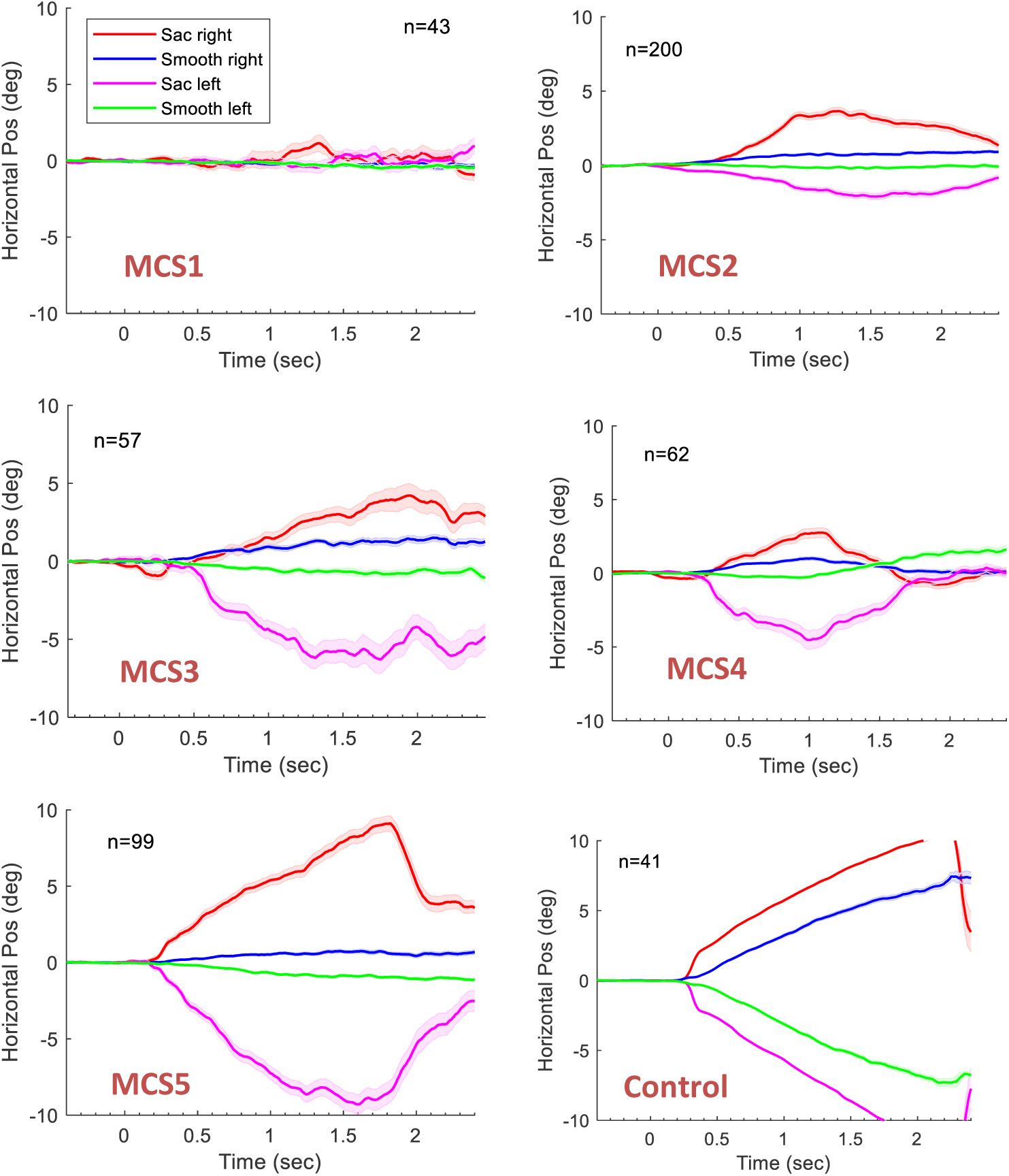
Pursuit horizontal eye movement in MCS patients, including individual data and control. The participants tracked a target that moved horizontally from the center (onset at time 0) left (negative) or right (positive H values) for 11°, changing direction every 6 movements (see the horizontal paradigm in the Methods section, Figure 1). The pursuit was analyzed in two ways: (1) including catchup saccades (saccadic pursuit, in red and magenta) and (2) excluding them from the traces (smooth pursuit, in blue and green). Individual data (all testing days) of MCS patients 1-5, and a group average of controls (n=5) are shown in separate plots, with identical axes for comparison. Error bars indicate the SE of the mean across trials (one movement per trial, 20-60 trials per participant). Note that compared with controls (lower right panel), the participants showed (1) a reduced and varied magnitude of movement, much below the target; (2) a large difference between the saccadic and smooth pursuit, with tracking done mostly by catchup saccades; and (3) a large asymmetry between sides in some participants (MCS1, MCS2).

We also examined the correlation between the clinical condition of the patient and the tracking performance. Figure 9 plots the correlation between the tracking velocity and the Lowenstein Communication Scale (LCS) obtained on each testing day for all 5 MCS patients. Importantly, we found a high correlation between these measures, primarily the saccadic pursuit velocity (R=∼0.8, see the Results, Fig. 9). Although the LCS clinical scale does contain a visual responsiveness item (1 of 5), the high correlation we found is remarkable. Apparently, the pattern of “fragmented tracking” via catchup saccades is the main characteristic property of the MCS patient’s condition and improves with recovery during the rehabilitation process. Our visual tracking paradigm could thus be used for assessing the patient’s condition.

**Figure 9.**
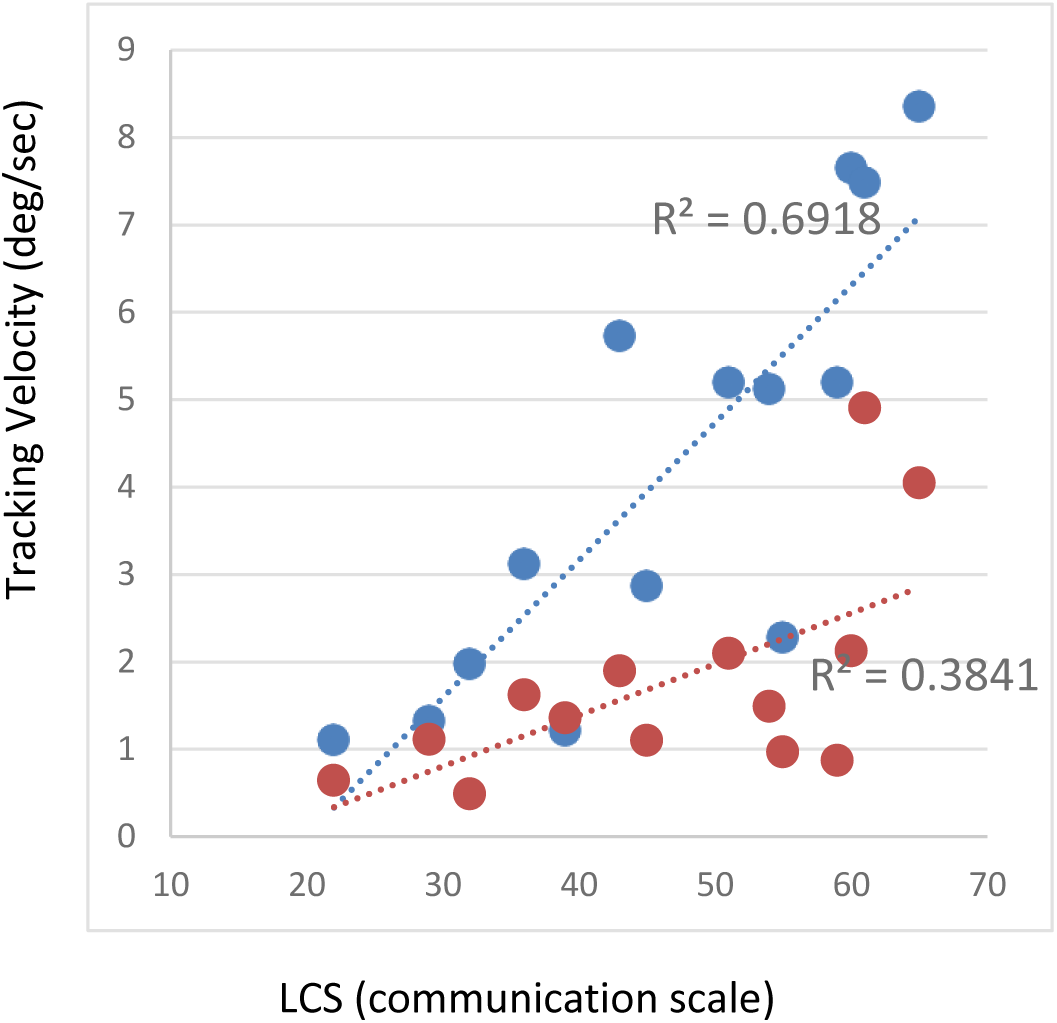
Correlation between the tracking velocity and the Lowenstein Communication Scale (LCS) for all 5 MCS patients, in each of their testing periods (3 numbers per patient, excluding 1). Blue – full pursuit, Orange – smooth pursuit (saccades excluded). The data are based on the horizontal traces of the Horizontal tracking experiments. Note the high and significant correlation for the full (saccadic) pursuit (R=∼0.83), with a somewhat lower correlation for the smooth pursuit data.

## Discussion

Oculomotor behavior is generally known as a good indicator of the severity and condition of TBI patients (Stubbs et al. 2019). In our study, we used a simple smooth pursuit paradigm using remote bedside eye-tracking technology that was found to produce a sensitive measure of traumatic brain injury. In the results of the first experiment with the standard linear pursuit movement, we found a difference between the TBI and the control groups in several measures of the pursuit quality. In the results of the second experiment with the pursuit movement under a visible occluder and an occluder assimilated into the background, the patients exhibited a larger deviance induced by the occluder.

### The oculomotor indices of TBI

We developed 6 oculomotor measures or indices to quantify the quality of eye movements during linear pursuit as a marker for TBI. These measures included the saccadic pursuit, tracking deviation under occlusion, initial tracking speed, initial saccade latency, pupil response, and vergence instability. The analysis of the indices revealed significant differences between the healthy and TBI groups (Figure 5). The FIM index was correlated with all the oculomotor indices with significance (R=∼0.4-0.8).

Beyond the observed deficits in smooth pursuit, our findings highlight a significant increase in interocular lag in TBI patients, particularly during horizontal tracking (Figures 4d, 5f). This suggests a disruption in binocular coordination, potentially reflecting an impairment in the vergence mechanisms or in the asymmetric neural control of both eyes. Similar disruptions in interocular coordination have been observed in macular degeneration (MD) patients, where smooth pursuit deficits are accompanied by increased interocular lag due to reliance on peripheral vision and impaired binocular integration (Shanidze et al. 2016). These parallels suggest that interocular lag may serve as a general marker of impaired sensorimotor integration across different neurological and visual conditions. Such disruptions may contribute to visual instability and difficulty in tracking moving objects in daily life, particularly in dynamic environments where precise binocular coordination is required.

Our findings indicate abnormal tracking behavior during occlusion in TBI patients. Unlike healthy controls, these patients failed to show the predictive ‘jump ahead’ eye movement that typically anticipates where the target will reappear after passing behind an occluder (Figure 3). A previous study by Diwakar et al.(Diwakar et al. 2015) used a virtual occluder (gap) with circular smooth pursuit and found increased tracking errors in chronic mild TBI patients specifically during target occlusion, but not during unoccluded tracking. Interestingly, they observed via MEG increased beta amplitude in the control group across several parietal regions, presumably reflecting the increased cognitive effort or attention required to make a predictive jump. Such cognitive effort should theoretically be reflected in increased pupil dilation. In our study, we found increased pupil dilation in the control group during simple tracking (Figure 4c). However, the bright occluders in our paradigm induced a strong pupil constriction response that complicates the interpretation of these findings.

The predictive ‘jump ahead’ effect observed in our controls is consistent with previous research demonstrating that smooth pursuit enhances motion prediction, allowing for accurate extrapolation of occluded trajectories (Kreyenmeier et al. 2022; Spering et al. 2011). The absence of this effect in TBI patients suggests not merely a failure to integrate velocity information, but also a breakdown in the internal forward model—a predictive process that allows the brain to estimate future sensory states based on motor commands and previous experiences (Molinari and Masciullo 2019; Wolpert and Miall 1996). This aligns with the notion that TBI, often involving diffuse axonal injury, affects higher-order cognitive control mechanisms responsible for predictive inference, rather than purely stimulus-driven tracking that forces TBI patients to rely on reactive tracking strategies instead of anticipatory ones.

### GCS versus Functional Measures in TBI Assessment

The widespread use of the Glasgow Coma Scale (GCS) as a measure of TBI severity, particularly in defining mild TBI (e.g., (Diwakar et al. 2015)), warrants critical examination. The relevance of the initial field-assessed injury severity to a patient’s subsequent condition raises important questions about using GCS for treatment decisions and prognosis. Our study found no correlation between GCS and FIM scores (p=0.8), despite strong correlations between FIM and various oculomotor measures (Figure 7). Similarly, none of our 6 oculomotor measures showed a significant correlation with GCS (see the Results, average p=0.59, ranging from 0.15 to 0.99). These findings suggest that cognitive indices reflecting the patients’ condition during rehabilitation, or functional measures like FIM, may be more appropriate than GCS for assessing TBI severity.

### Comparison to previous studies

Although previous research has extensively studied eye movements in TBI, most studies have focused on mild TBI cases classified by GCS scores rather than functional status (Hunfalvay et al. 2019, 2020, 2021; Maruta, Lee, et al. 2010; Maruta, Suh, et al. 2010; Suh et al. 2006; Zahid et al. 2017). Our study demonstrates that simpler experimental methods can provide clinically relevant measures. Previous studies used more complex approaches such as circular tracking (Diwakar et al. 2015; Hunfalvay et al. 2019, 2020, 2021; Maruta et al. 2014, 2016, 2017, 2018; Maruta, Lee, et al. 2010; Maruta, Suh, et al. 2010; Suh et al. 2006), target under occlusion (Diwakar et al. 2015), or sustained sinusoidal tracking (Hunfalvay et al. 2020). In contrast, we found that a simple repeated linear tracking task with 0.5-second pauses between segments was sufficient to reveal significant group differences. This simplified approach reduces patient fatigue and data complexity, making it more suitable for clinical applications. Additionally, our use of portable eye tracking without head restraints enabled comfortable bedside testing in rehabilitation settings—a crucial factor for clinical adoption. Finally, whereas previous studies mainly identified group differences between TBI patients and controls without establishing links to functional outcomes, our study demonstrates strong correlations between eye-tracking measures and FIM scores, directly connecting oculomotor dysfunction to functional independence.

### Eye Tracking with DOC Patients

To extend our findings, we applied the same tracking paradigm to a preliminary sample of 5 DOC patients, whose conditions were more severe than typical TBI cases. Our findings diverged slightly from previous studies in visual pursuit within DOC. Unlike prior research, which primarily reported smooth pursuit without fine-tuned computerized eye tracking, we observed that visual pursuit was absent in our small sample of Unresponsive Wakefulness syndrome (UWS) patients. Moreover, in minimally conscious state (MCS) patients, we noted a distinct pattern of fragmented visual tracking, characterized by saccadic, rather than smooth pursuit. This fragmented saccadic pursuit, with sequential catch-up saccades, appears to reflect a fundamental characteristic of DOC, potentially mirroring fragmented perceptual awareness. Notably, saccadic pursuit velocity strongly correlated with Lowenstein Communication Scale (LCS) scores across testing days, highlighting its potential as a clinical marker. This correlation underscores the fragmented tracking behavior as a key indicator of MCS, which improves with rehabilitation and recovery. Thus, this eye-tracking paradigm holds promise as a clinical tool for assessing DOC patient conditions, tracking recovery progress, and, in specific cases, identifying injury characteristics such as lateral asymmetry. Overall, our findings suggest that fragmented saccadic tracking serves as a marker of DOC, potentially aiding in distinguishing it from TBI and assessing recovery. Further research is needed to validate catch-up saccades as a diagnostic marker for awareness impairment and for assessing the severity of disruption in DOC.

### Potential for future diagnosis and prognosis of TBI

Our results suggest that a simple and short bedside recording of a linear visual tracking examination of a few minutes could serve as an effective tool for providing measures of inefficient brain functioning after traumatic brain injury. Our detailed assessment of short and effortless visual tracking revealed markers that are very sensitive to the patient’s condition and therefore can be useful for clinical evaluation. These markers can be used in the future to reveal injury-specific deficits. For example, a horizontal interocular deviation could indicate an injury to the 6th nerve in the eye opposite the affected hemisphere. Moreover, these measures, combined perhaps with machine learning techniques, could be used to develop a good prognosis and evaluation of the efficacy of treatments. Future research should focus on combining reliable measures to develop practical, fast, and affordable user-friendly tools for doctors and professionals.

Beyond TBI assessment, our findings highlight the diverse nature of oculomotor impairments following traumatic head injury, which affect the initial tracking speed and latency, vergence instability, pupil-linked arousal, as well as the ability to predict and adjust to dynamic visual stimuli. These deficits may contribute to everyday challenges, such as reading or navigating dynamic environments. Our results suggest that fundamental eye movement behaviors—such as smooth pursuit and saccadic adjustments—could serve as useful indicators of these impairments. This underscores the potential value of incorporating eye-tracking measures into clinical assessments to better capture and understand both motor and cognitive dysfunction in TBI patients. Such an approach may help monitor recovery and refine rehabilitation strategies in a practical and objective manner.

## Author contributions

SS, YSB, and EG designed the experiments. SS, YS, and KC collected the data in the main study (TBI), and EG, OK, and LA collected the data in the DOC study. YSB developed the experimental tool, and YSB and OK developed the data analysis code. SS and YSB analyzed the data in the main study, and EG, OK, and YSB analyzed the data in the DOC study. SS and YSB wrote the paper, and OK and YS reviewed it.

## Data availability

The experimental datasets generated during the current study will be available from the corresponding author upon request.

