## Supplemental Tables for "Bedside Assessment of Visual Tracking in Traumatic Brain Injury: Comparing Simple and Predictive Paradigms Using Multiple Oculomotor Markers"

Supplementary

| | DOC | Age | Sex | Etiology | LCS | Time post onset (months) |
|---|---|---|---|---|---|---|
| VS1 | UWS | 38 | M | MVA | 15 | 3 |
| VS2 | UWS | 30 | M | SAH (AVM), Multiple Infarctions | 15 | 1 |
| VS3 | UWS--MCS | 44 | M | Fall | 27 | 4 |
| MCS1 | MCS--TBI | 57 | M | Fall | 13 | 2 |
| MCS2 | MCS | 55 | F | AVM repair | 29 | 4 |
| MCS3 | UWS--MCS | 21 | M | MVA | 22 | 3 |
| MCS4 | MCS--TBI | 27 | M | Assult | 32 | 2 |
| MCS5 | MCS | 33 | M | MVA | 43 | 2 |
| MCS6 | MCS--TBI | 66 | F | ICH | 30 | 2 |
| MCS9 | MCS | 24 | F | MVA | 41 | 2 |
| MCS11 | UWS | 69 | M | MVA | 33 | 2 |

**Table S1:** Patients' demographic and clinical features. MVA-motor vehicle accident, ICH-intracerebral hemorrhage, SAH- subarachnoid hemorrhage, and AVM- arteriovenous malformation. The LCS (Lowenstein Communication Scale) was measured around the time of each day of testing: data from the first day of examination are shown here.

| | |
|---|---|
| DOC | Disorder of Consciousness |
| UWS | Unresponsive Wakefulness Syndrome |
| MCS | Minimally Conscious State |
| Hc | Saccadic pursuit |
| H0 | Smooth pursuit |
| OKN | Optokinetic nystagmus |
| MVA | Motor vehicle accident |
| ICH | Intracereberal hemorrhage |
| SAH | Subarachnoid hemorrhage |
| AVM | Arteriovenous malformation |
| LCS | Lowenstein communication scale |

Table S2: Abbreviation List
